# Systemic infection of SARS-CoV-2 in free ranging Leopard (*Panthera pardus fusca*) in India

**DOI:** 10.1101/2022.01.11.475327

**Authors:** Sonalika Mahajan, Mathesh Karikalan, Vishal Chander, Abhijit M. Pawde, G Saikumar, M Semmaran, P Sree Lakshmi, Megha Sharma, Sukdeb Nandi, Karam Pal Singh, Vivek Kumar Gupta, Raj Kumar Singh, Gaurav Kumar Sharma

## Abstract

We report patho-morphological and virological characterization of SARS-CoV-2 in naturally infected, free ranging Indian Leopard (*Panthera pardus fusca*). Whole genome sequence analysis confirmed infection of Delta variant of SARS-CoV-2, possibly spill over from humans, but the case was detected when infection level had dropped significantly in human population. This report underlines the need for intensive screening of wild animals for keeping track of the virus evolution and development of carrier status of SARS-CoV-2 among wildlife species.

---

Currently there are increasing reports of the natural infection of severe acute respiratory syndrome coronavirus (SARS-CoV-2) in various domesticated and captive wild species [Reviewed in 1-2]. Both *in-silico* [3] and experimental infections of [4] SARS-CoV-2 have been studied in number of species that detailed out the immunogenicity and pathogenesis of the virus in these species. However, infection in species other than humans is considered as spill over infection and so far there are no reports of transmission of this virus among animal population on large scale.

Although intensive investigations were carried out, only three cases of natural infection in Asiatic lions could be detected in India [5–6]. India experienced huge second wave with surge in cases from the month of March 2021 with peak reaching in the month of April when more than 18 million cases were reported in a month (Fig 1a; https://www.mohfw.gov.in/). Surge in the cases were attributed to spread of highly infectious Delta variant (Pango lineage B.1.617.2) of SARS-CoV-2 and lack of herd immunity [7]. With surge of infection and vaccination coverage, herd immunity in the population improved and number of cases decreased. With drop in human infections in India, fresh cases were not found in animals during random screening. During the months of September and October-21, cases in India dropped to less than 10,000/day, however, in the second week of October-21 one Indian leopard cub (*Panthera pardus fusca)* was found dead and upon investigation was found positive for SARS-CoV-2. In this report we are presenting the gross and histopathological findings, presence of viral antigen in tissues and genetic character of the virus.

**Figure 1:**
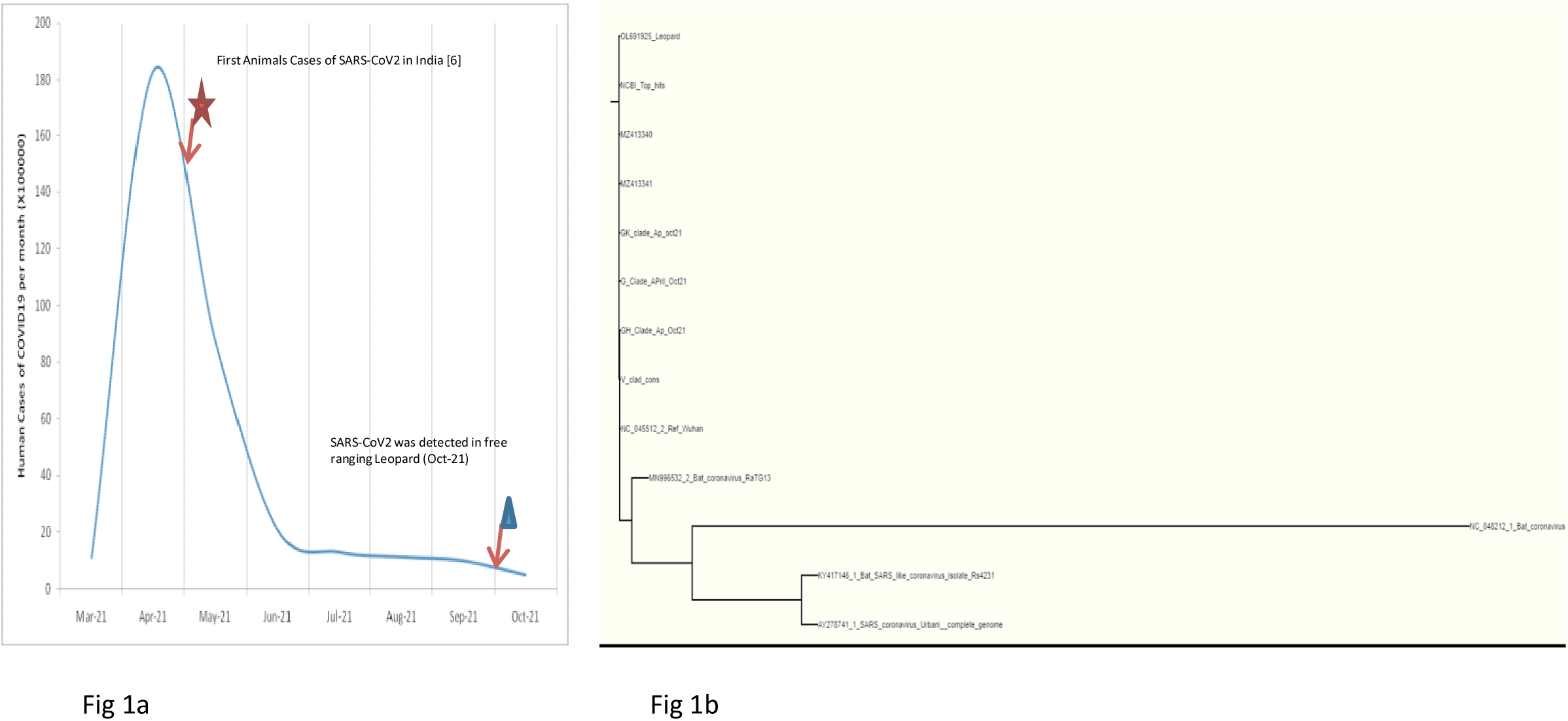
**1a.** Number of COVID-19 cases in humans and detection of animal infections in India **1b.** Phylogenetic analysis of SARS-CoV-2 detected in an Indian leopard. Whole-genome phylogeny with consensus sequences of other clades available in the GISAID database revealed the generated sequence clustered with the G-clad (B.1.617.2 pango lineage).

The carcass of male leopard cub of approximately of 1 year age was recovered during routine combing in the agricultural field of Mojipur village of Social Forestry, Bijnor range of Uttar Pradesh State (29.372442°N and 78.135849°E), located approximately 160 kilometer from national capital of New Delhi. Bijnor Social Forestry Division has approximately 14,400 hectare of reserve forest located in the foot hills of the Himalaya. About 70% of the inhabited land is having sugarcane cultivation which favours the habitation of wild herbivores and carnivores in this area.

The carcass was submitted to the ICAR-Indian Veterinary Research Institute for necropsy examination. Necropsy was conducted following COVID-19 protocol wearing PPE kit *etc*. Animal carcass and the waste material generated were incinerated and area was disinfected using 2% hypochlorite solution.

Necropsy examination revealed piercing wounds on both the sides of the ventral neck region (canine teeth marks of prey animal), subcutaneous contusions and haemorrhages on neck and cranium suggesting case of infighting (Supplementary Fig 1). Additionally, consolidation of both the lungs (Supplementary Fig 1), and severe vascular changes like congestion and haemorrhages in the visceral organs were observed. Rectal and nasopharyngeal swabs and tissue samples from various organs were collected and subjected to routine laboratory investigations. Nasopharyngeal swab was found positive for SARS-CoV-2 by RT-PCR using COVISure Viral detection kit (Genetixbiotech, India). Hence, all collected organs (Supplementary table 1) were once again tested and results were confirmed by the generating partial spike protein gene sequence using Sanger’s method as described previously [6].

Samples were also screened and ruled out for presence of any other pathogens including haemoprotozoan or rickettsial parasites, canine distemper virus, Feline panleukopenia virus, Feline herpes-virus and Feline calicivirus, and highly pathogenic Avian Influenza (H5 and H7 as described previously [6].

Positive nasopharyngeal swab was subjected to virus isolation in Vero cells as described previously and blindly passaged for 7 passages and observed for obvious cytopathic effects (CPE). Presence of virus was confirmed by RT-PCR and immuno-staining with SARS-CoV-2 positive serum followed by detection by FITC labeled anti-human secondary antibodies (Real gene, Germany).

We generated whole genome sequence directly from the nasal swab specimen through outsourcing to Eurofins as described previously [6] and submitted to NCBI with GenBank accession ID OL691925. The SRA project was submitted with accession number PRJNA786974. The BLAST tool indicated >99.9% identity with SARS-CoV-2 sequence (OK189615-OK189617) collected from Kerala state of India during the month of September 2021. Compared to the Wuhan-Hu-1 sequence (NC-045512), 40 amino acid substitutions were observed of which 3 were present in the 5’UTR and 2 were in 3’UTR (Supplementary table 2). Six amino acid substations *viz.*Thr19Arg, Asn99Lys, Leu452Arg, Thr478Lys, Asp614Gly, and Pro681Arg were observed in the spike protein. Spike protein sequence of virus from leopard showed very high resemblance with that of sequences of Delta variant and with the sequences generated in previous study from infected Asiatic lions from Jaipur and Etawah in India [6]. However, two amino acid substitutions (Thr77Lys and Asp142Gly) were observed when compared to genome sequences from infected Asiatic lions of Tamil Nadu [5]. Although very few animals have been found infected by SARS-CoV-2, high resemblance of spike protein sequences with that of human Delta variant suggests possible spill over infection and no major genetic changes in the spike protein upon species cross over.

To elucidate the phylogenetic and temporal analysis, we downloaded representative sequences from GISAID database of each clade of SARS-CoV-2 samples collected from India during the period of March 2021 to Oct 2021. Consensus sequences of each clade was generated by aligning using ClustalW tool in MEGA X version 10.1 followed by consensus generation in EMBOSS server. Phylogeny was constructed using NGphylogeny server as described previously [6]. The leopard sample clustered into common SARS-CoV-2 G-clade (GISAID classification) or Delta variant (B.1.617.2 Pango lineage) (Fig 1b). The leopard sequence was closely matching with the prevailing Delta mutant in the area suggesting possible spill over infection in the animal.

Brain, spleen, lymph node and lungs specimens were found positive for SARS-CoV-2 with Ct values of E-gene ranging from 27.5-31.6 (Supplementary table 1). Histopathologically, the lung revealed diffuse areas of consolidation, hemorrhages, pneumocyte hyperplasia, septal thickening and perivascular infiltration of mononuclear cells (Fig 2). Severe vascular changes were also noticed in the heart, brain, liver and kidneys. The spleen and lymph nodes showed mild depletion of lymphoid follicles. Viral antigen was demonstrated in the lungs, brain and spleen by immunohistochemistry using human anti-SARS-CoV-2 hyperimmune sera and goat antihuman IgG-HRPO antibodies (Real Gene, Germany) as per previously described protocol [8]. Abundant viral antigen was noticed in the septal lining cells, alveolar macrophages, endothelial cells of pulmonary vessels and bronchiolar epithelial cells (Fig 2). Similarly, viral antigen was detected in the lymphoid cells and macrophages of spleen and glial cells and capillary endothelium of brain (Fig 2).

**Figure 2:**
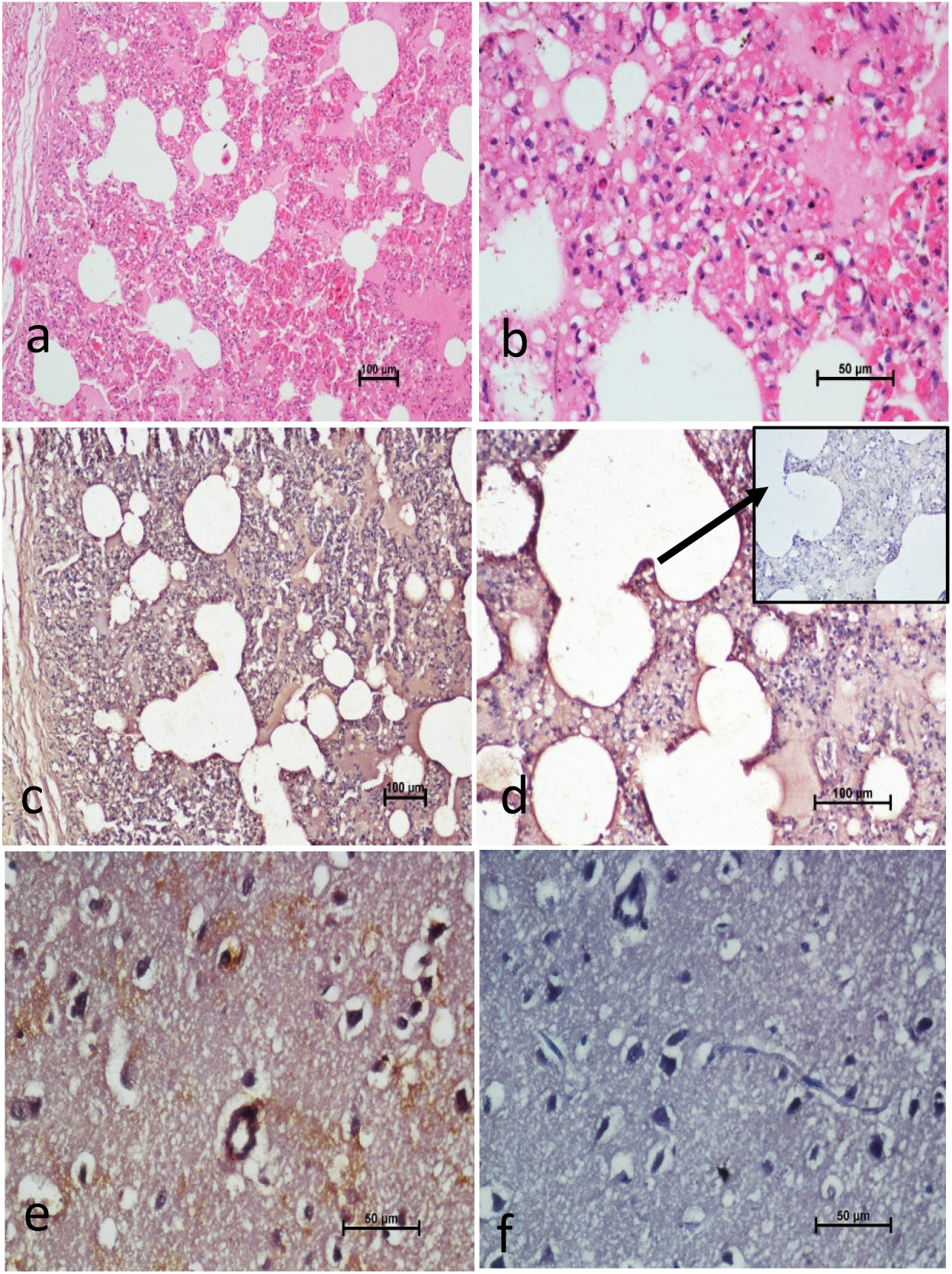
2a. The lung showing diffuse inflammatory changes like haemorrhageand congestion and thickening of alveolar septum. HE x100 2b. Higher magnification fig. a showing alveolar septal thickening and infiltration of mononuclear cells and pneumocyte hyperplasia. HE x400 2c. The lung showing strong immune reactivity to SARS-COV-2 in the alveolar septal cells and inflammatory cells. IHC DAB x100. 2d. Higher magnification fig. c showing strong immune reactivity to SARS-COV-2 in the alveolar septal cells and inflammatory cells. IHC DAB x200. Inset: Antibody control, IHC DAB x200 2e. The cerebrum showing strong immune reactivity to SARS-COV-2 in the capillary endotheial cells, glial cells and neuron. IHC DAB x400. 2f. The cerebrum showing absence immune reactivity to SARS-COV-2 (antibody control). IHC DAB x400.

Several natural infections in minks and humans and experimental infection of SARS-CoV-2 in laboratory animals showed similar lung histopathological lesions as observed in the present case [10–12].

Though the necropsy findings suggested traumatic injuries as the immediate cause of death; detection of virus in various tissues indicates systemic SARS-CoV-2 infection of the animal prior to sudden death due to trauma inflicted by another carnivore. Presence of virus in the brain section suggests infection of central nervous system in wild felid which has previously been demonstrated in humans only [9, 13]. Previously, one SARS-CoV-2 infected captive Asiatic lion died at Chennai, India (unpublished), however, necropsy examination could not be conducted to ascertain the exact cause and co-morbidities.

The forest range in Bijnor is protected but shares border with human habitation. On an average, 10-15 leopards die of either natural cause or diseases and there is no alarming increase in the mortality in the area hence it seems to be a focal case of SARS-CoV-2 infection. Leopards are relatively less shy when compared to other wild felids which may provide opportunity for spill over events in this species. Most of the studies in animals were reported either in domestic or captive wild animals with a history of human contact. Detection of SARS-CoV-2 in free ranging leopard when incidences of human COVID19 have dropped to significantly lower level (Fig 1a) underlines the necessity to intensify the screening and check for development of carrier status in wild felids.

## About the authors

Dr Sonalika is Scientist at ICAR-Indian Veterinary Research Institute, Izatnagar Bareilly UP India. Her primary research interest includes diagnosis and vaccine development against emerging viruses of animal origin.

## Data availability

Annotated complete genome sequences are available in NCBI (GenBank accession: OL691925). Raw sequencing data are available at NCBI BioSample (Accession; PRJNA786974).

## SUPPLEMENTAL MATERIAL

**Supplementary Table 1;** qPCR Ct values of SARS-CoV-2 positive samples

**Supplementary Table 2**: Comparison of nucleotide and amino acid mutations in SARS-CoV-2 isolated from Leopard sequence in comparison to the Reference Wuhan Strain (NC_045512.2)

**Supplementary Figure 1:** Animal carcass examination for external wounds and Gross examination of Lungs and brain.

## ACKNOWLEDGMENTS

Authors are thankful to the Bijnor Forest Division for submitting the carcass for necropsy examination. We are also thankful to ICAR, NASF and DST-SERB for providing funds and infrastructure to take up the work.

## Conflict of interest

Authors disclose no conflict of interest with anyone

## Notes

### Competing Interest Statement

The authors have declared no competing interest.

