## supplementary tables for "Systemic infection of SARS-CoV-2 in free ranging Leopard (*Panthera pardus fusca*) in India"

**Supplementary Table 1;** qPCR Ct values of SARS-CoV-2 positive samples

| **S No** | **Samples** | **Gene 1**  **(E Gene)** | **Gene 2**  **(RdRP)** |
| --- | --- | --- | --- |
| 1 | Nasal swab | 27.5 | 29.6 |
| 2 | Rectal swab | 29.7 | 31.2 |
| 3 | Brain | 30.3 | 32.8 |
| 4 | Spleen | 29.4 | 29.8 |
| 5 | Lymph node | 31.6 | 33.5 |
| 6 | Lung | 27.9 | 29.9 |

Supplementary Table 2: Comparison of nucleotide and amino acid mutations in SARS-CoV-2 isolated from Leopard sequence in comparison to the Reference Wuhan Strain (NC_045512.2)

| **Position** | **Reference Allele** | **Alternate Allele** | **Read Depth** | **Mapping quality** | **Gene Name** | **Protein Change** |
| --- | --- | --- | --- | --- | --- | --- |
| 201 | T | G | 1009 | 60 | Intergenic (CHR_START-ORF1ab) | - |
| 210 | G | T | 1056 | 60 | Intergenic (CHR_START-ORF1ab) | - |
| 241 | C | T | 1075 | 60 | Intergenic (CHR_START-ORF1ab) | - |
| 774 | C | T | 1104 | 60 | ORF1ab | Thr170Ile |
| 2937 | C | T | 6732 | 60 | ORF1ab | Thr891Ile |
| 3037 | C | T | 7861 | 60 | ORF1ab | Phe924Phe |
| 5184 | C | T | 7999 | 60 | ORF1ab | Pro1640Leu |
| 5544 | C | T | 7979 | 60 | ORF1ab | Thr1760Ile |
| 5584 | A | G | 7205 | 60 | ORF1ab | Thr1773Thr |
| 6449 | C | T | 4675 | 60 | ORF1ab | Leu2062Phe |
| 9891 | C | T | 5593 | 60 | ORF1ab | Ala3209Val |
| 10776 | C | A | 318 | 60 | ORF1ab | Pro3504His |
| 11418 | T | C | 7126 | 60 | ORF1ab | Val3718Ala |
| 12946 | T | C | 7980 | 60 | ORF1ab | Tyr4227Tyr |
| 13904 | A | G | 6541 | 60 | ORF1ab | Asp4547Gly |
| 14408 | C | T | 697 | 60 | ORF1ab | Pro4715Leu |
| 15451 | G | A | 5025 | 60 | ORF1ab | Gly5063Ser |
| 16466 | C | T | 7978 | 60 | ORF1ab | Pro5401Leu |
| 18176 | C | T | 7967 | 60 | ORF1ab | Pro5971Leu |
| 20262 | A | G | 8000 | 60 | ORF1ab | Leu6666Leu |
| 21618 | C | G | 7312 | 60 | S | Thr19Arg |
| 21859 | C | A | 122 | 60 | S | Asn99Lys |
| 22917 | T | G | 1122 | 60 | S | Leu452Arg |
| 22995 | C | A | 1093 | 60 | S | Thr478Lys |
| 23403 | A | G | 7885 | 60 | S | Asp614Gly |
| 23604 | C | G | 7967 | 60 | S | Pro681Arg |
| 25469 | C | T | 7679 | 60 | ORF3a | Ser26Leu |
| 25562 | A | G | 7991 | 60 | ORF3a | Gln57Arg |
| 26767 | T | C | 8000 | 60 | M | Ile82Thr |
| 27046 | C | T | 7998 | 60 | M | Thr175Met |
| 27638 | T | C | 7958 | 60 | ORF7a | Val82Ala |
| 27739 | C | T | 7055 | 60 | ORF7a | Leu116Phe |
| 27752 | C | T | 6555 | 60 | ORF7a | Thr120Ile |
| 28249 | A | T | 11 | 60 | ORF8 | Asp119Val |
| 28253 | C | A | 55 | 60 | ORF8 | Phe120Leu |
| 28881 | G | T | 7964 | 60 | N | Arg203Met |
| 29402 | G | T | 7992 | 60 | N | Asp377Tyr |
| 29541 | C | T | 7998 | 60 | Intergenic (N-ORF10) | - |
| 29742 | G | T | 7716 | 60 | Intergenic (ORF10-CHR_END) | - |
