## Supplementary figure for "Systemic infection of SARS-CoV-2 in free ranging Leopard (*Panthera pardus fusca*) in India"

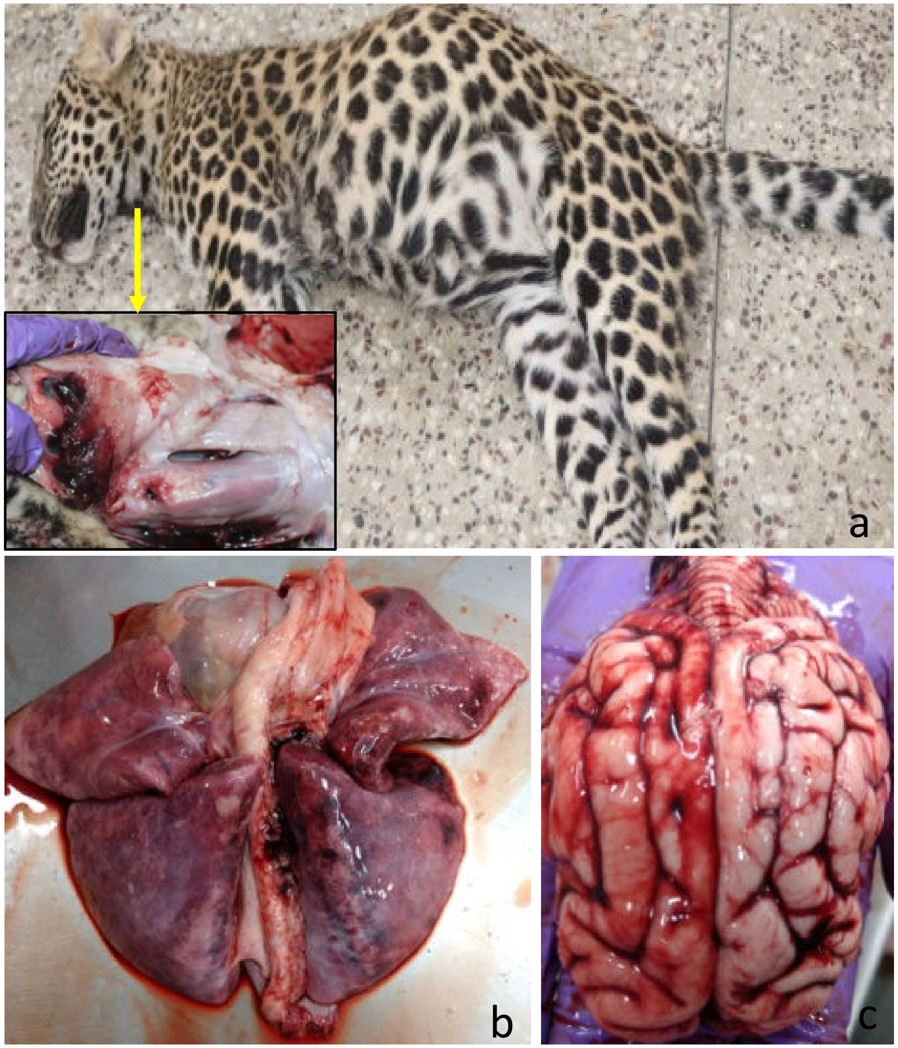


Supplementary Figure 1:

a. Carcass of male leopard cub, Inset: Ventral neck having piercing wounds (teeth marks of prey animal)

b. The lung showing diffuse consolidation with red and grey patches

c. The brain showing severely congested blood vessels
